# Reverse engineering of a pathogenic antibody reveals the molecular mechanism of vaccine-induced immune thrombotic thrombocytopenia

**DOI:** 10.1101/2023.06.30.547300

**Authors:** Daniil G. Ivanov, Nikola Ivetic, Yi Du, Son N. Nguyen, S. Hung Le, Daniel Favre, Ishac Nazy, Igor A. Kaltashov

## Abstract

The massive COVID-19 vaccine roll-out campaign illuminated a range of rare side effects, the most dangerous of which – vaccine-induced immune thrombotic thrombocytopenia (VITT) – is caused by adenoviral (Ad)-vectored vaccines. VITT occurrence had been linked to production of pathogenic antibodies that recognize an endogenous chemokine, platelet factor 4 (PF4). Mass spectrometry (MS)-based evaluation of the ensemble of anti-PF4 antibodies obtained from a VITT patient’s blood indicates that its major component is a monoclonal antibody. Structural characterization of this antibody reveals several unusual characteristics, such as the presence of an *N*-glycan in the Fab segment and high density of acidic amino acid residues in the CDR regions. A recombinant version of this antibody (RVT1) was generated by transient expression in mammalian cells based on the newly determined sequence. It captures the key properties of VITT antibodies, such as their ability to activate platelets in a PF4-dependent fashion. Homology modeling of the Fab segment reveals a well-defined polyanionic paratope, and the docking studies indicate that the polycationic segment of PF4 readily accommodates two Fab segments, cross-linking the antibodies to yield polymerized immune complexes. Their existence was verified with native MS by detecting assemblies as large as (RVT1)_3_(PF4)_2_, pointing out at FcγRIIa-mediated platelet activation as the molecular mechanism underlying VITT clinical manifestations. In addition to high PF4 affinity, RVT1 readily binds other polycationic targets, indicating a polyreactive nature of this antibody. This surprising polyspecificity not only sheds light on VITT etiology, but also opens up a range of opportunities to manage this pathology.

**Significance Statement:** Vaccine-induced immune thrombotic thrombocytopenia (VITT) is a dangerous side effect of adenoviral-vectored vaccines that is linked to the emergence of autoantibodies recognizing platelet factor 4 (PF4). We have engineered a recombinant VITT antibody by sequencing a VITT patient-derived anti-PF4 monoclonal antibody that causes platelet activation and triggers thrombosis. This antibody was used to characterize architecture of the pathogenic immune complexes with a combination of biophysical and computational approaches, revealing the molecular mechanism of VITT. The results of this work demonstrate the critical role of electrostatics in PF4 recognition by the pathogenic antibody and the polyspecificity of the latter. Availability of the engineered VITT antibody will be invaluable for future studies aiming at understanding the general mechanistic features of autoimmune pathologies.

## Introduction

Vaccine-induced immune thrombotic thrombocytopenia (VITT) is a rare but extremely dangerous (and frequently fatal) side effect of adenoviral (Ad) vectored vaccines that came to light during their massive roll-out and use in the COVID-19 pandemic (1). VITT typically occurs within 4-28 days of vaccination and has a distinct clinical profile (thromboses at unusual sites, such as cerebral venous sinus thrombosis (2, 3) in combination with thrombocytopenia) and a high mortality rate (4). Despite the low incidence rate (ranging from 3 to 36 cases per million doses for ChAdOx1 and three- to four-fold lower for human Ad-vectored vaccines (5, 6)), VITT became one of the major factors contributing to the vaccine hesitancy phenomenon and undermining the global vaccination effort during the pandemic (7-9). Although all VITT cases reported so far had been linked to COVID-19 vaccines, the close association of this thrombopathy with a specific delivery vector raises a specter of other Ad-vectored vaccines being able to trigger VITT, an alarming prospect given the popularity of this platform (10).

VITT presentation is strikingly similar to another thrombopathy, heparin-induced thrombocytopenia (HIT) (11). Furthermore, VITT mirrors HIT in that it is associated with the emergence of pathogenic antibodies recognizing a small chemokine, platelet factor 4 (PF4, also referred to as CXCL4) (6, 12). While this naturally leads to a suggestion that the anti-PF4 antibodies are critical players in FcγRIIa-mediated platelet activation in VITT just as they are involved in HIT pathogenesis (13), there is an important distinction. VITT development does not require heparin, a critical element of the pathogenic HIT-related immune complexes, where it not only enables accommodation of multiple PF4 molecules within a single immune complex (14), but also causes a conformational change within PF4 (15). The latter is sufficient to expose a neo-epitope on the PF4 surface (15), thereby making this endogenous protein a legitimate “not self” target for the immune system.

The critical importance of heparin in the etiology of HIT and its notable irrelevance to VITT despite the striking similarity of the two pathologies prompted several groups to propose scenarios where heparin’s dual role of (*i*) the scaffold of the immune complex assembly and (*ii*) the effector of PF4 conformational change is assumed by other anionic biopolymers, such as polyanionic impurities (16), the viral DNA - the Ad vector cargo (17), or indeed the surface of the Ad capsid (18). However, mapping the VITT antibodies’ epitopes on the PF4 surface revealed their significant overlap with the heparin-binding sites (19), lending credibility to the notion of the immune complex formation via cross-linking of antibodies by polyvalent (tetrameric) protein antigens and unassisted by any polyanionic scaffolds/effectors. The realization that the epitope repertoire is limited also led to a suggestion that the VITT antibodies’ clonality is significantly restricted; indeed, the oligo- (and in some cases mono-) clonal nature of VITT antibodies was subsequently reported (20, 21).

The restricted clonality of VITT antibodies indicates that their formation does not follow the classic B-cell maturation mechanism, but instead proceeds via rapid expansion of a limited number of B-clones (21). This raises a question of what allows this process to bypass the elaborate “self/not-self” selection mechanisms. Furthermore, the specific architecture of the immune complexes capable of triggering platelet activation and eventually precipitating VITT remains elusive. While the ability of an Ad capsid to adsorb PF4 on its surface has been demonstrated (18), implying that the former may act as a scaffold for assembling VITT immune complexes, the feasibility of the platelet-activating complexes formation in the absence of Ad capsids has also been demonstrated (22). Until this dichotomy is addressed, and the complete picture of the molecular mechanism underlying VITT pathology becomes available, rational development of effective prophylactic and therapeutic strategies targeting VITT will remain challenging at best. In fact, the on-going efforts to develop the “VITT risk-free” Ad vectors by designing capsids lacking PF4-binding elements (23) may prove misguided should no template be needed for the pathogenic immune complex formation (19, 22, 24). Furthermore, the very identity of PF4 as the “genuine” antigen for the pathogenic VITT antibodies was questioned recently following the demonstration of their ability to bind another chemokine, CXCL7 (25).

Our work aims at closing these gaps by performing a detailed structural analysis (complete amino acid sequencing) of a monoclonal anti-PF4 antibody extracted from the blood of a VITT patient and creating its recombinant copy using transient expression. This reverse-engineered antibody (designated RVT1) binds PF4 and activates platelets in a PF4-dependent fashion. Using native MS, we demonstrate that PF4 acts as a multi-valent antigen by cross-linking RVT1 molecules to form polymerized immune complexes, the likely agents of FcγRIIa-mediated platelet activation. The availability of the VITT antibody sequence also allows homology modeling to be used to build its 3D model, which reveals a well-defined anionic paratope. This not only provides a strong support to the notion of the antibody/antigen interactions being driven primarily by electrostatic interactions, but also explains how the cationic PF4 tetramers polymerize the antibodies, giving rise to immune complexes capable of FcγRIIa-mediated platelet activation, without the need for polyanionic scaffolds. The electrostatic nature of the VITT antibody/PF4 interaction also hints at the possibility of this antibody’s recognizing and binding to other polycationic proteins. Indeed, native MS provides conclusive evidence of RVT1 cross-reactivity towards other polybasic targets, such as protamine. Lastly, an unusual structural feature of the VITT antibody (also captured in RVT1) is the presence of an *N*-glycan within the Fab region. While this glycan does not appear essential for the antigen binding, its presence may explain the “self-reacting” clone survival and selection, as has been recently proposed for a range of other autoimmune disorders (26, 27).

## Results

### Reverse engineering of the anti-PF4 VITT-associated monoclonal antibody

Our study relies on availability of a molecularly defined model that captures all the essential structural and functional properties of the pathogenic anti-PF4 VITT antibodies (28). There is a growing consensus that these antibodies have a significantly restricted clonality (20, 21), making it possible to obtain exhaustive amino acid sequence(s) information. The anti-PF4 antibodies were extracted from the peripheral blood of a patient who developed multiple thrombotic complications following ChAdOx1-S vaccination and was subsequently diagnosed with VITT based on the results of the PF4-enhanced heparin-independent EIA assay as well as serotonine-release assays according to the current clinical guidance (29) (see ***Supplementary Methods*** for detail). Intact-mass MS analysis of the IgG molecules affinity-precipitated on the PF4-conjugated beads reveals a bimodal mass distribution showing a distinct contribution of a well-defined ionic signal representing a set of glycoforms of a single antibody (**Figure S1** in ***Supplementary Material***). The LC-MS analysis of the light chain, and the Fc/2 and Fd segments of the heavy chain (generated by the proteolytic cleavage of the antibody below the hinge region followed by disulfide reduction within all fragments) confirms its monoclonality. PNGase F treatment of the antibody resulted in no changes of the light chain mass, while introducing a mass shift of the Fc/2 segment to 23,792±3 Da, thereby confirming the absence of N-glycans within the light chain and the presence of a heterogeneous and immature bi-antennary *N*-glycan of the complex type within the Fc/2 segment, typical of IgG molecules (**Figure 1**). In addition, a mass shift of the Fd segment to 25,422±3Da following enzymatic de-*N*-glycosylation reveals the presence of a fully mature bi-antennary *N*-glycan of the complex type within the Fd segment, a less common site of *N*-glycosylation within γ-immunoglobulins (30). The occurrence of *N*-glycans within the Fab regions of the IgG molecules had been noted in the past to be correlated with the emergence of monoclonal antibodies in several auto-immune disorders (26, 27, 31). Our subsequent efforts were directed towards de novo sequencing of this monoclonal antibody and localizing the two *N*-glycosylation sites.

**Figure 1.**
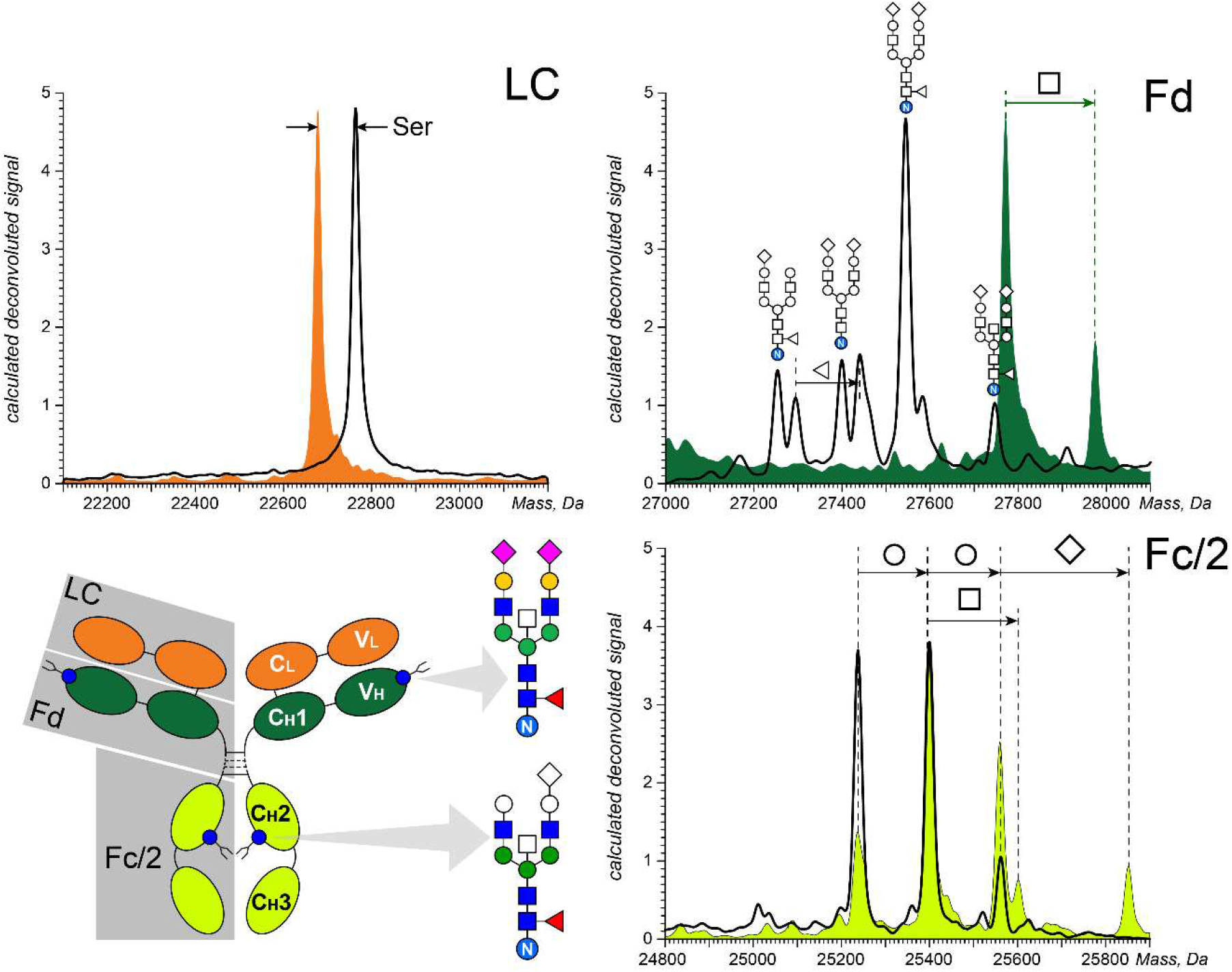
Deconvoluted mass spectra of the light chain, and the Fd and Fc/2 segments of the heavy chain of VITT mAb (color-filled curves) and RVT1 (black curves). The schematic representation of the two N-glycans (***bottom left***) shows the color-filled base structures (corresponding to the lowest-mass glycoforms of the VITT mAb detected by MS); the higher-mass glycoforms are shown by addition of colorless monosaccharide symbols.

De novo sequencing of the anti-PF4 monoclonal antibody was carried out using peptide mapping and LC-MS/MS analysis of the proteolytic fragments. Two complementary peptide maps were produced using trypsin and chymotrypsin to enable alignment of the fragment peptides. Since the traces of polyclonal antibodies were not removed from the antibody sample prior to peptide mapping, their proteolytic fragments were also present in the digest, and an intensity-based criterion was used to filter out peptides not representing the monoclonal component. The resulting sequences of both light and heavy chains of the monoclonal antibody (VITT mAb) are presented in **Figure 2**. Residues 108-213 of the light chain line up with the IGLC3 gene (32), confirming that the light chain belongs to the λ-type. The V_L_ domain aligns best with the IGLV3-21*02 gene, displaying only five mutations in the LCDR regions (two and three for LCDR1 and LCDR3, respectively), two mutations in the framework 3 region and one deletion at the N-terminus. The paucity of mutations is also evident upon aligning the heavy chain sequence with the IGH genes, showing only six deviations from the IGHV2-26*02 sequence in the entire V_H_ segment excluding HCDR3. Two of these mutations fall outside of the CDR1/CDR2 segments, and both are localized within the framework region 3. Importantly, one of them (K75N) creates an *N*-glycosylation motif (***N***XT), which was confirmed with LC-MS/MS to be the site of *N*-glycosylation within the Fab segment of the antibody. While the HCDR3 gene sequences are not available in the IMGT database (32), we note a significant similarity of the sequence determined in our work with those reported earlier for a set of VITT patients (20). Indeed, the highly acidic GLEDAFD motif evident in the HCDR3 region characterized in this work bears significant resemblance to the sequences (G/N)LED(A/V/T)FD reported by Wang et al. (20). This similarity is noteworthy, since the HCDR3 is the most variable region among all CDRs (being a product of recombination of three genes, V_H_, D and J). The sequences of the C_H_1, C_H_2 and C_H_3 domains, as well as that of the hinge region, are identical to those of the IGHG2 gene, allowing the monoclonal antibody to be assigned to the IgG2 subclass.

**Figure 2.**
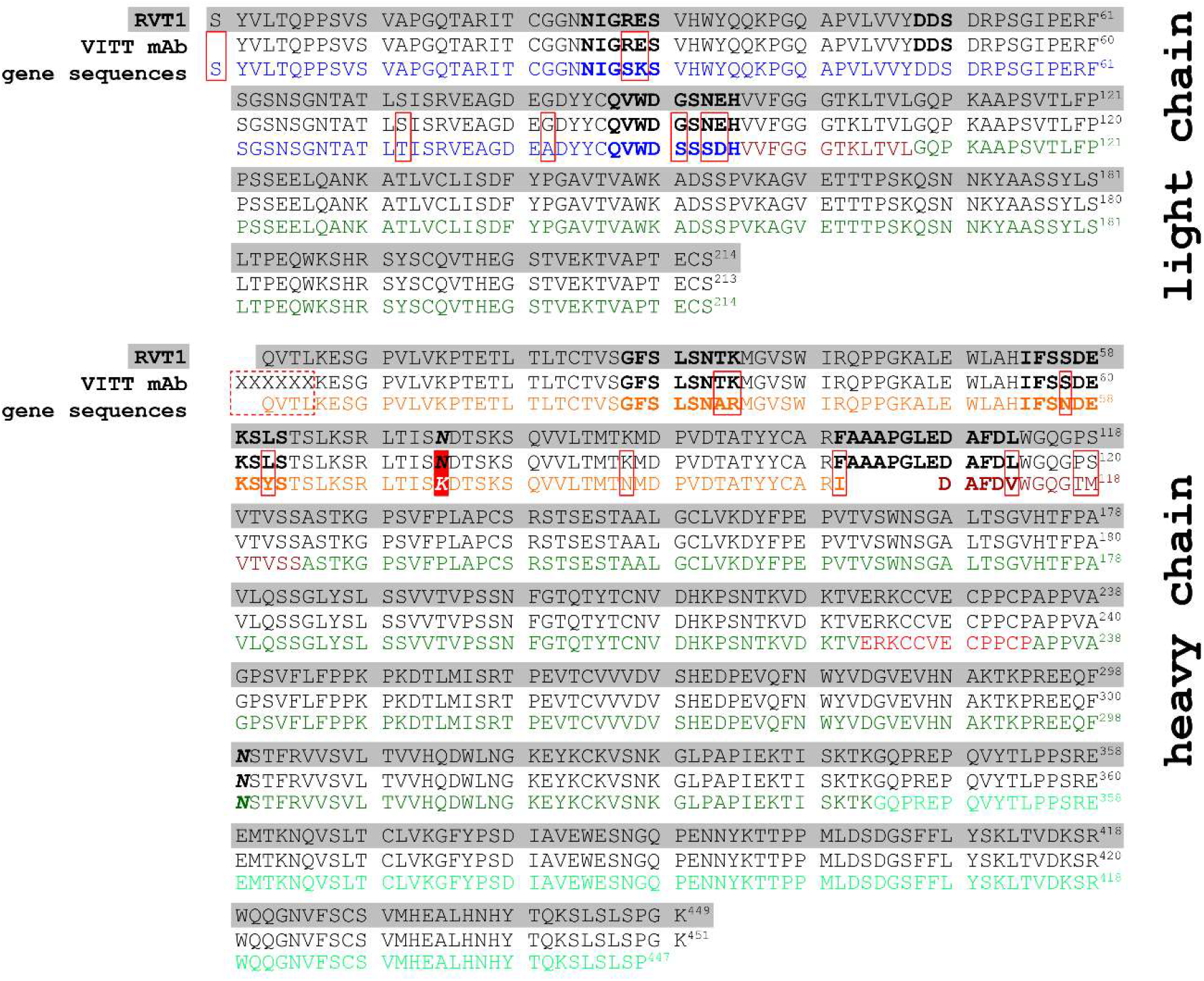
Amino acid sequences of light (***top***) and heavy (***bottom***) chains of RVT1, VITT mAb and the most closely matching germline gene sequences (the VITT mAb sequence was constructed on the basis of peptide mapping and LC/MS/MS analysis of the monoclonal anti-PF4 antibody extracted from the VITT patient’s plasma – see the ***Supplementary Material*** for more detail). The underlined residue is missing in the mAb sequence, and the residues in parentheses have not been positively confirmed. The CDR regions are typed in bold, and the N-glycosylation sites within the V_H_ and C_H2_ domains are typed in bold italic. The colored sequences represent the following genes: IGLV3-21*02 (blue), IGLJ2/IGLJ3 (maroon) and IGLC3 (green) for the light chain; IGHV2-26*02 (orange), IGHJ (maroon), and IGHG2 (C_H_1, C_H_2 and C_H_3 domains -green, hinge region domain - red) for the heavy chain. Note that RVT1 authentication revealed the post-production loss of the C-terminal lysine residue and the N-terminal glutamine conversion to pyro-glutamate (see the ***Supplementary Material*** for more detail).

The VITT mAb sequence determined by peptide mapping and LC-MS/MS was used to construct a recombinant copy of this antibody. The de novo determined sequence of the light chain was appended with an N-terminal serine residue to make the production more robust, and the undetermined six-residue N-terminal sequence of the heavy chain was replaced with a QVTL tetrapeptide (to match the N-terminal part of the IGHV2-26*02 gene. The recombinant copy of the VITT monoclonal antibody was generated using transient expression via co-transfection of two vectors containing heavy and light chains as shown in **Figure 2**. This antibody, referred to as RVT1, was expressed in HEK293T cells, purified on a protein A resin and extensively characterized prior to being used in antigen-binding and platelet-activation properties testing. Characterization of transiently expressed RVT1 revealed no post-translational modifications (PTMs) within its light chain and two PTMs within the heavy chain (conversion of the N-terminal Gln residue to *pyro*Glu and removal of the C-terminal Lys residue). Both PTMs are common among mAbs and are innocuous (neither the antibody stability nor its antigen-binding properties are compromised by these PTMs (33)).

### Antigen binding and platelet-activating properties of VITT RVT1

The initial evaluation of RVT1 biological properties was carried out by measuring its affinity to the presumed antigen (PF4) and the ability to activate platelets. The PF4/RVT1 association strength was measured with SPR on a Protein A-conjugated chip, yielding an apparent K_d_ value of 330±80 nM (**Figure S2** in **Supplementary Material**). The behavior of RVT1 vis-à-vis platelet activation was probed with the standard and PF4-enhanced SRA, and found to follow the pattern that is common to VITT-related pathogenic antibodies (19). No platelet activation was observed in the standard SRA for RVT1, while the following PF4 dose-dependent platelet activation responses were recorded in the PF4-enhanced SRA: 2% release at 10 µg/mL PF4, 20% at 25 µg/mL, and 53% at 50 µg/mL. In all experiments, platelet activation levels were inhibited following the addition of IV.3, a monoclonal anti-CD32 antibody that blocks the FcγRIIa platelet antibody receptor, confirming that the RVT1-triggered platelet activation is indeed mediated by FcγRIIa. The sub-μM PF4 affinity of RVT1 and its ability to activate platelets in the PF4-dependent fashion while being negative in the standard SRA confirms that RVT1 captures the essential properties of the pathogenic anti-PF4 antibodies extracted from the VITT patient’s blood and, therefore, can be used as a reliable model antibody to study the molecular mechanism of VITT pathogenesis.

### Characterization of the RVT1/PF4 immune complexes architecture

The availability of a recombinant antibody that captures the essential biological properties of the anti-PF4 antibodies extracted from a VITT patient’s blood (and has a structure near-identical to that of the monoclonal component of the anti-PF4 IgG ensemble) enables mechanistic studies of the disease pathogenesis. Our work focused on characterizing molecular architecture of immune complexes whose formation is precipitated by PF4. While characterization of immune complexes containing multiple antigen and antibody molecules remains challenging due to their large size and heterogeneity, native MS has been recently demonstrated to be capable of providing meaningful information on such macromolecular assemblies (34-36). High molar excess of PF4 over RVT1 in aqueous solutions at near-physiological pH and ionic strength gives rise to mass spectra dominated by an abundant ionic signal corresponding to the 2:1 antigen/antibody complexes, but shows no signal indicative of the presence of large immune complexes in solution (the blue trace in **Figure 3**). However, adjusting the antibody/antigen ratio in solution to a nearly-equimolar level (3.5 μM each) results in a dramatic change in the appearance of the mass spectrum, which now displays abundant ionic signals representing PF4-polymerized antibodies (RVT1_2_PF4, RVT1_2_PF4_2_ and RVT1_3_PF4_2_) in addition to the uncomplexed antibody (the red trace in **Figure 3** and **Figure S3** in ***Supplementary Material***). The presence of the abundant ionic signal at *m/z* above 12,000 suggests that even larger immune complexes (exceeding 500 kDa) are populated in solution, but these remain unassigned due to the lack of resolved/clearly discernable spectral features. It is noteworthy that formation of the RVT1*_n_*PF4*_n-1_* and RVT1*_n_*PF4*_n_* complexes detected by native MS does not require the presence of obligatory polyanionic scaffolds (such as heparin in HIT). The preferred *n*:*(n-1)* antibody/antigen stoichiometry of the observed immune complexes provides a clear indication that it is the cross-linking of RVT1 molecules by PF4 that gives rise to these assemblies. At the same time, high antigen/antibody ratios inhibit the polymerization process by capping both Fab segments of nearly all RVT1 molecules with PF4, thereby preventing further growth of the complex, as is observed by native MS in RVT1/PF4 mixtures with large molar excess of the latter (*vide supra*).

**Figure 3.**
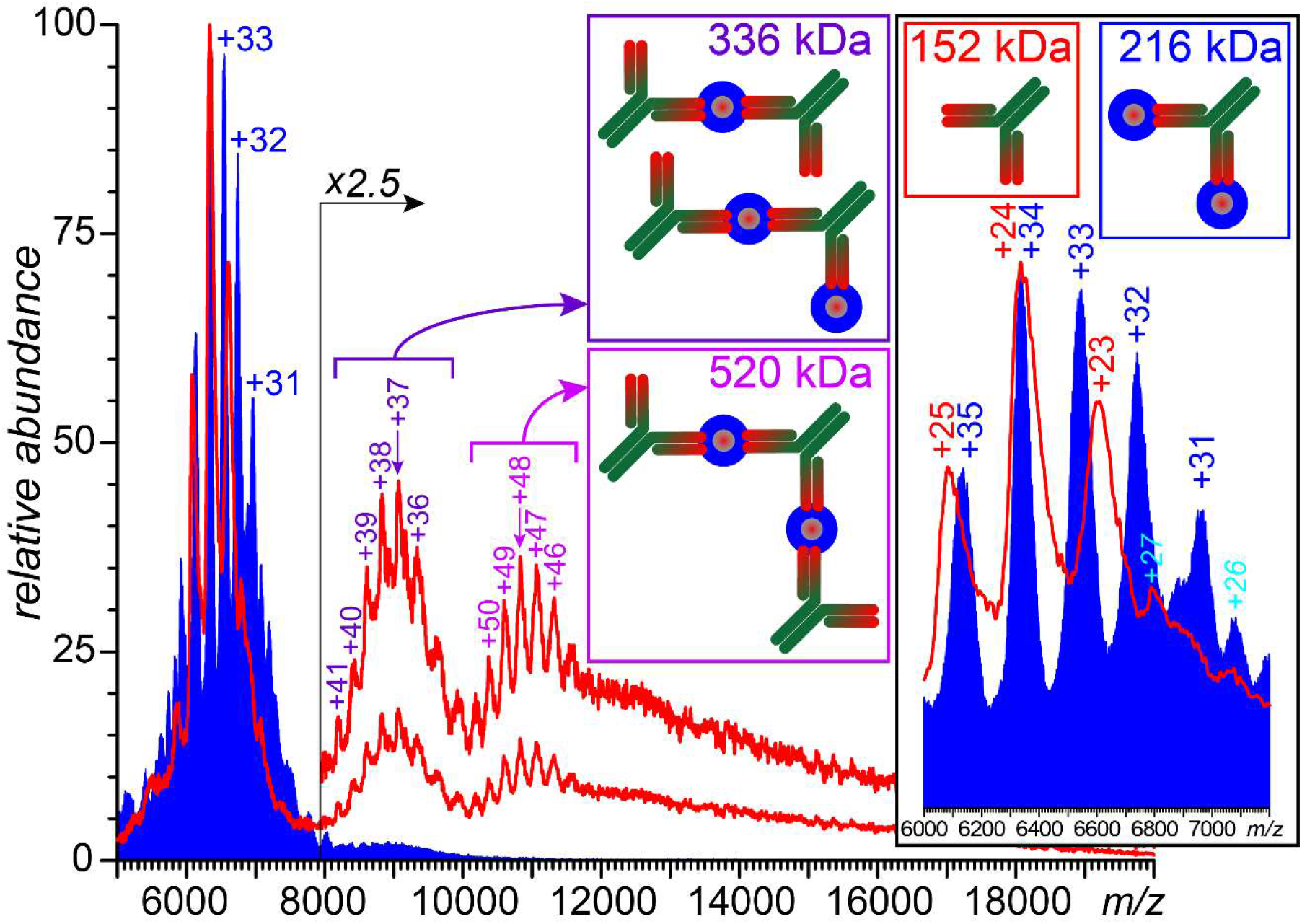
Native mass spectra of RVT1/PF4 mixtures acquired with 1:5 (blue) and 1:1 (red) antibody/antigen molar ratios in solution.

### Formation and growth of the pathogenic immune complexes are driven by the epitope/paratope charge complementarity

All native MS measurements described in the preceding paragraph were carried out using protein solutions maintained at the physiological ionic strength. The increase of the solution ionic strength results in a monotonic decrease of the abundance of the antibody/antigen complexes, and their signal completely disappears from the mass spectra above 1 M of salt (see **Figure S4** in **Supplementary Material**). Such a dramatic dependence of the antibody/antigen interaction efficiency on the solution ionic strength is a clear indication of the electrostatic nature of this association (37). To explore the role of electrostatics in PF4 recognition by *RVT1* in greater detail, a 3D model of its Fab segment was generated using homology modeling. The resulting energy-minimized structure maintains a classic IgG fold despite the presence of several mutations within the framework regions both in the light and heavy chains (**Figure 4**). Calculation of the electrostatic potential distribution for this structure reveals a well-defined patch of the negative charge, which is formed by five of the six CDR regions (**Figure 4**). Specifically, 6 aspartic acid and 2 glutamic acid residues within these segments are arranged circularly, and the resulting ring forms a paratope that should be able to act as an electrostatic trap for positively-charged epitopes – such as the heparin-binding regions of PF4. Therefore, it is not surprising that docking of the Fab segment to a tetrameric PF4 shows a strong preference of this polyanionic paratope for the positively charged regions of the antigen’s surface. The best-scoring epitopes identified by docking contain residues 17-25, 45-50 and 60-70 that form positively charged segments on the PF4 surface (see **Figure S5** in ***Supplementary Material***). These segments includes all 7 residues of the protein identified by Huynh et al. as critical for the VITT antibodies binding based on alanine scanning (19). However, it should be noted that reasonably good docking scores were also obtained for other segments of the protein, as long as they were located within the “positive charge belt” circumscribing the PF4 tetramer, indicating that the antibody interaction with its target is rather promiscuous (see **Figure S6** in ***Supplementary Material***).

**Figure 4.**
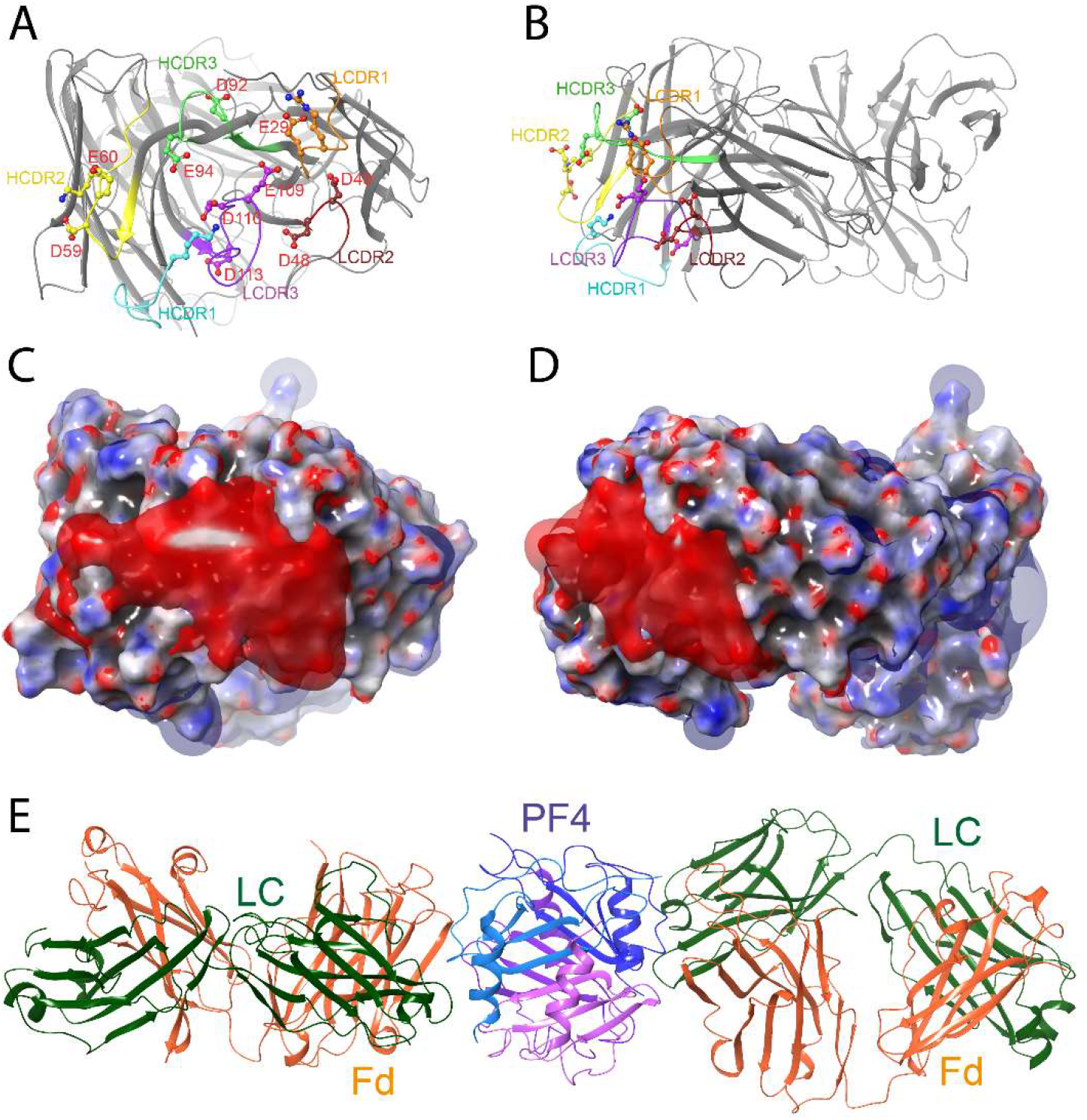
***A-B***: Structure of *RVT1* Fab segment produced by homology modeling represented by the front (**A**) and side (**B**) views of Fab showing all relevant CDR loops and acidic residues forming the polyanionic paratope. The space-filled model of Fab and the electrostatic iso-potential surfaces (3*k*T/*e*) are shown in **C** and **D** using the same orientations. Panel **E** shows the energy-minimized structure of two RVT1 Fabs cross-linking by a single PF4 tetramer produced by molecular docking.

Since the polypeptide chains are arranged tetrahedrally within the PF4 tetramer, there are two identical patches on its surface that present the best-scoring RVT1 docking site. This allows a single PF4 tetramer to readily accommodate two Fab segments in the trans configuration, as confirmed by docking of an RVT1 Fab fragment to the Fab·PF4 complex generated by the initial docking step. The resulting Fab·PF4·Fab complex (**Figure 4E**) provides an insight into the antibody cross-linking by a bivalent antigen, although one must be mindful of the promiscuous nature of the PF4 interaction with the RVT1 Fab fragment. The latter may give rise to immune complexes with not only alternative epitopes, but also different binding stoichiometry. Indeed, our docking studies indicate that a single PF4 tetramer can accommodate up to three RVT1 Fab fragments, and extension of these fragments to the full-length antibodies shows that there are no steric clashes that may preclude formation of such “dendritic” immune complexes (RVT1)_3_·PF4 (see **Figure S7** in ***Supplementary Material***).

### Electrostatics as the driving force in antigen recognition and RVT1 cross-reactivity

The defining role of electrostatic interactions in PF4 recognition by RVT1 determined by native MS is rationalized by the molecular modeling studies. However, the latter also reveals a certain degree of promiscuity in the paratope/epitope interactions (*vide supra*). This raises a question of whether other polycationic proteins (or low-pI proteins with well-defined positively charged surface segments) can also be recognized by the polyanionic paratopes of RVT1. The question of RVT1 cross-reactivity was approached in our work by using protamine as a model polycation. This choice was predicated on the fact that the polypeptides from the protamine family exhibit extremely high density of the positive charge (due to the incorporation of multiple poly-Arg segments in their sequences), while at the same time lacking any well-defined structure in solution (38). Analysis of the RVT1/protamine mixture with native MS provides clear evidence that a single antibody molecule readily accommodates up to two protamine polypeptides at the physiological ionic strength (**Figure S8** in ***Supplementary Material***). No larger complexes (e.g., two antibody molecules cross-linked by a single protamine polypeptide chain) were detected in these measurements, consistent with the small size of these polypeptide, which precludes accommodation of multiple antibodies by a single antigen. Above and beyond revealing the cross-reactivity of *RVT1*, these measurements pose an intriguing question of whether PF4 is in fact the primary target of the “anti-PF4” VITT antibodies, rather than their unfortunate “bystander” victim that happened to display the characteristics of an ideal target – well-defined regions of high-density positive charge.

## Discussion

The majority of mechanistic studies of VITT-associated thrombotic events continue to rely on the patients-derived heterogenous PF4-binding serum (19, 39). The scarcity of this clinical material and its limited availability to researchers outside of a few specialized hematology laboratories unfortunately limit both the scope of such studies and the pace of progress in this field. The complete structural characterization of the monoclonal component of the anti-PF4 antibodies extracted from a VITT patient’s blood presented in this work enables construction of a recombinant form of this antibody (RVT1) that correctly captures the key biological properties of the VITT antibodies. It can be produced in milligram quantities in facilities capable of transient expression of recombinant immunoglobulins, thereby lending itself as not only an extremely useful and unique vehicle for mechanistic studies of VITT pathogenesis at the molecular level, but also a readily available and affordable one. The model VITT antibody RVT1 designed by reverse engineering of the monoclonal component of the anti-PF4 VITT antibodies activates platelets in the heparin-independent, PF4-sensitive fashion – the key biological property that is common to all VITT-associated anti-PF4 antibodies and in fact is currently the basis for VITT diagnosis in clinical laboratories (40). Furthermore, RVT1 structure bears significant resemblance to the sequences of the LCDR3 and HCDR3 segments of oligoclonal antibodies from several unrelated VITT patients (20), which are the most variable antibody segments. All this makes the RVT1 molecule designed and tested in our work a truly paradigmatic VITT antibody model that is clearly indispensable in the studies of the molecular mechanism of this pathology in general, rather than representing only one specific clinical case.

Both native MS analysis and molecular modeling studies of RVT1 provide a strong indication that the properties shared by all VITT-associated anti-PF4 antibodies are due to the high concentration of acidic amino acid residues within the paratope (**Figure 4**). The resulting high-density negative charge basin localized within the antigen-binding region allows this antibody to interact effectively with the heparin-binding sites of PF4, where high-density positive charge is localized (36, 41). As PF4 in its native tetrameric state exhibits a contiguous polycationic region – a high positive charge-density equatorial belt circumscribing the entire protein – its recognition by the VITT antibody does not require any conformational changes within PF4. This is a major point of distinction between VITT and HIT, where heparin binding to the protein triggers a conformational change that allosterically creates a neoepitope at a remote site, thereby enabling the PF4 recognition by pathogenic antibodies (15). Above and beyond being capable of binding PF4, RVT1 can be readily cross-linked by this protein (as confirmed by native MS and molecular modeling), giving rise to macromolecular assemblies capable of clustering FcγRIIa receptors on the platelet surface, a key step triggering their activation (**Figure 5**). This is another important distinction of VITT, which – unlike HIT (14), and contrary to the earlier suggestions (16-18, 23) – does not require a polyanionic scaffold to enable antigen clustering as a prerequisite for assembling large pathogenic immune complexes. Indeed, the electrostatic nature of the epitope/paratope interaction and the presence of the extended polycationic segment on the PF4 surface allow it to cross-link at least two VITT antibodies, as confirmed by native MS measurements where abundant complexes as large as (RVT1)_3_(PF4)_2_ have been identified (**Figure 4**).

**Figure 5.**
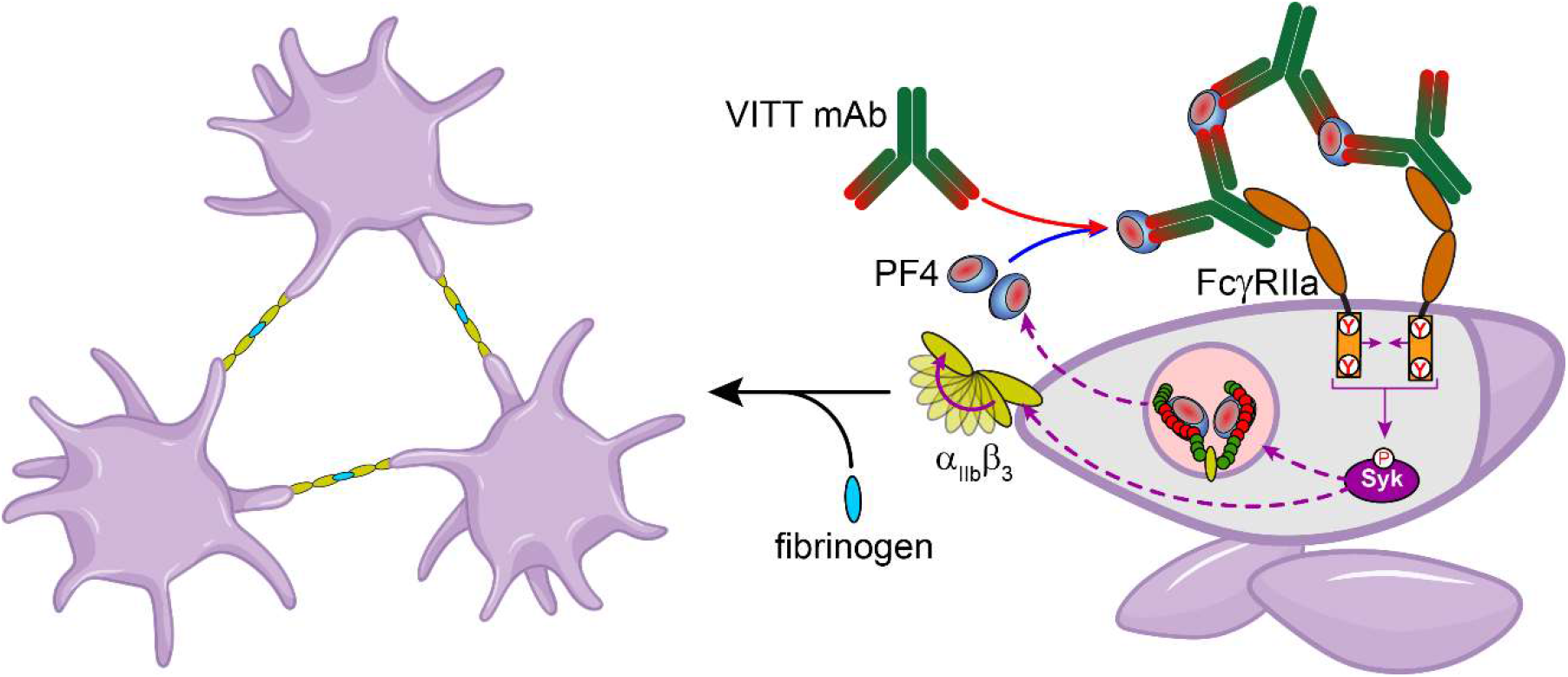
A schematic representation of the mechanism of platelet activation by PF4-polymerized VITT antibodies.

The role of electrostatics as a major determinant of the VITT antibody specificity may seem surprising, but is certainly not unique, as evidenced by the anti-DNA antibodies in systemic lupus erythematosus (42). At the same time, the dominance of electrostatics in defining the paratope/epitope association brings to the fore the question of cross-reactivity of VITT antibodies. Indeed, there is a large number of proteins displaying well-defined patches of positive charge (as suggested by the heparin interactome size (43)), which naturally lend themselves as viable candidates vis-à-vis the putative target pool of VITT antibodies. In fact, a recent observation points out at the ability of VITT antibodies to associate with another polycationic chemokine, NAP2 (also referred to as CXCL7) in addition to PF4 (25). While PF4 (CXCL4) and CXCL7 bear significant structural resemblance to each other, our native MS measurements provide conclusive evidence that even structureless polycations – such as protamine – are readily recognized by RVT1, a testament to their remarkable degree of promiscuity. Another likely – yet untested – target of VITT antibodies that is relevant in the thrombogenesis setting is neutrophil elastase, an important player in NETosis (44). The paradoxical conclusion that despite having severely restricted clonality, VITT antibodies possess broad specificity also invites scrutiny vis-à-vis the very labeling of these antibodies as “anti-PF4.” Indeed, it is not unreasonable to suggest that the actual target of these antibodies remains unidentified, while the PF4 molecules are accidental bystander victims due to their polycationic character, and their recognition by and association with the VITT antibodies become the defining element of the disease pathogenesis due to the PF4 levels in circulation spiking as a result of inter alia immune system activation in the wake of vaccination (45, 46).

While the identity of the original antigen of VITT antibodies remains unknown, the strongly anionic character of the paratope may provide some clues vis-à-vis viable candidates, *e.g*. positively charged segments of the vector capsid, such as the heparan sulfate-binding element(s) of the Ad fiber shaft. At the same time, the remarkable closeness of the VITT mAb structure to the germline gene sequences and the paucity of mutations (**Figure 2**) suggests this is not a fully mature/optimized antibody, and that an alternative selection mechanism should be invoked to explain its massive production in a VITT setting. Indeed, one of the very few mutations evident within the VITT mAb amino acid sequence converts a lysine residue (followed by aspartic acid and threonine residues in the sequence) to asparagine, thereby creating a novel *N*-glycosylation site, which is found to bear a fully mature biantennary *N*-glycan of a complex type within VITT mAb (a feature preserved in RVT1). While the presence of this structural feature in the variable region of an IgG molecule has been traditionally thought of as very rare, recent estimates suggest that over 15% of the entire IgG ensemble in the serum may be *N*-glycosylated (30). Furthermore, this structural feature appears to be common to many autoimmune disorders involving production of pathogenic antibodies from a limited number of clones, including rheumatoid arthritis (26), primary Sjögren’s syndrome (27), myasthenia gravis (47), and several others (48). In most cases the novel *N*-glycan within the Fab region adds little-to-none as far as the antigen binding affinity (contrary to an earlier view stipulating a positive contribution (49)). This leads to an argument that the commonality of *N*-glycosylation within the variable domains in autoimmune diseases hints at its role in providing a selective advantage to the clone by enabling alternative forms of selection to the corresponding B-cells (27). While the details of this alternative selection mechanism remain elusive (50), its existence explains both the proliferation of an apparently immature clone that produces monoclonal VITT antibodies, and the cross-reactivity of the latter. Indeed, a significant fraction of the nascent B-cells express autoreactive antibodies (51). While most of them are eliminated during the maturation process, a rogue early mutation generating an *N*-glycosylation site within the Fab region would not only allow this clone to bypass the elaborate “self/not self” selection machinery, but in fact make it the predominant one, thereby triggering production of monoclonal (but polyspecific) self-reacting antibodies.

While the molecular mechanism and the proposed etiology of VITT presented in this work imply that an a priori prediction of this this side effect’s occurrence or even identification of the risk groups is nearly impossible, certain properties of VITT antibodies exemplified by our model RVT1 molecule may inform the search for novel effective therapeutic strategies. While most current practices rely on therapeutic plasma exchange, high-dose intravenous immunoglobulin injections, and corticosteroids (52), the cross-reactivity of RVT1 and the polyanionic nature of the paratope region suggest that relatively small polycationic substances may lend themselves as safe and potent inhibitors of the VITT antibody/PF4 interactions. The high positive charge density within such substances will allow them to act as the VITT antibody paratope binders/blockers, while their small physical size will prevent them from cross-linking the antibodies, thereby eliminating the specter of creating large immune complexes capable of platelet activation via the FcγRIIa-mediated route.

The happenstance nature of VITT occurrence and its association with a specific vector (the reports of this side effect outside of the Ad-vectored vaccines remain extremely rare (53)) suggest that its relevance extends far beyond COVID-19 vaccines. Indeed, the gargantuan scale of the vaccination campaign during the recent pandemic and its initial localization within countries with developed health care systems illuminated the problem that is likely to be inherent to other vaccines using the same or similar vector(s), which otherwise would have remained undetected for a long time. One particularly serious concern here is the popularity of this delivery platform in novel and effective vaccines that primarily target diseases in developing countries (*e.g*., Ebola, Zika, HIV and malaria (10)). The diagnostic capabilities in many of these localities may be inadequate or lacking altogether, thereby presenting a challenge vis-à-vis timely diagnosis and treatment of this side effect. This problem can be solved only by extensive and concerted efforts of researchers and health care practitioners addressing all aspects of this pathology, and VITT vigilance is certainly warranted despite the COVID-19 pandemic slowly receding into the history.

## Materials and Methods

### Study participant criteria and the VITT anti-PF4 antibody purification

The plasma sample used in this study was collected from a patient who had been administered the ChAdOx1-S vaccine and subsequently developed VITT. The diagnosis was made according to the currently accepted criteria based on the positive results of commercial PF4-enhanced EIA (Immucor, OD ≥ 0.45) and PF4-enhanced serotonin release assay (SRA ≥ 20% ^14^C-serotonin release) analysis. This study was approved by the Hamilton Integrated Research Ethics Board (HiREB). The entire IgG ensemble was purified from the patient’s plasma using Protein G-conjugated gravity column (Cytiva, Marlborough, MA) and loaded onto the streptavidin beads conjugated with biotynilated recombinant PF4. After the elution of the anti-PF4 antibodies from the beads, sample was buffer exchanged into the PBS and subsequently tested using the PF4-enhanced SRA analysis.

### Liquid chromatography/mass spectrometry (LC/MS) and de novo sequencing of the anti-PF4 antibodies

Following the intact-mass analysis of the purified anti-PF4 antibodies using a Synapt G2 HDMS (Waters Corp., Milford, MA) hybrid quadrupole/time-of-flight mass spectrometer, the proteins were digested with IdeZ and reduced with dithiothreitol. The resulting fragments (the light chain, as well as the Fc/2 and Fd segments of the heavy chain) were analyzed with LC/MS using a SolariX 7 (Bruker Daltonics, Billerica, MA) Fourier-transform ion cyclotron resonance mass spectrometer. PNGase F (NEB, Ipswitch, MA) was used for de-*N*-glycosylation. The complete sequencing and *N*-glycans localization of the VITT mAb were carried out using peptide mapping with two orthogonal proteases (trypsin and chymotrypsin) with and without prior de-*N*-glycosylation. The proteolytic fragments were analyzed by LC/MS/MS using Orbitrap Fusion (Thermo, San Jose, CA) mass spectrometer coupled to Easy-nLC 1000 nano-UHPLC system (C18 column). The initial data processing was carried out with a PEAKS Studio xPro (Bioinformatics Solutions, Waterloo, ON) software suit followed by manual inspection of all relevant MS/MS datasets. A comprehensive technical report detailing this procedure is available on *bioRxiv* (54).

### Recombinant expression of RVT1

Production of RVT1 was carried out by Sino Biological (Wayne, PA) in HEK293T cells co-transfected with two plasmids carrying synthetic genes using the transient expression technology. The produced antibodies were initially authenticated using gel electrophoresis, followed by a thorough MS-based analysis using the same protocol that had been employed for characterization and sequencing of the VITT mAb (*vide supra*).

### Native MS analysis of the polyvalent complexes

RVT1 association with the recombinant form of human PF4 and UPS-grade protamine (Millipore-Sigma, St. Louis, MO) were characterized using a Synapt G2 HDMS (Waters Corp., Milford, MA) hybrid quadrupole/time-of-flight mass spectrometer. All measurements were carried out in 150 mM ammonium acetate, pH 6.8, unless specified otherwise.

### Functional platelet activation assays

The platelet-activating capability of RVT1 were measured with standard and PF4-enhanced SRAs following the commonly accepted protocols (55). Plasma-derived platelets were radiolabeled with ^14^C-serotonin, and platelets were incubated with RVT1 both in the absence and in the presence of PF4. Platelet activation was monitored by measuring measure the release of ^14^C-serotonin using a scintillation counter (Packard Bell, Topcount).

### Surface-plasmon resonance (SPR)

The SPR measurements of RVT1/PF4 binding were carried out using a Biacore T200 (Cytiva) device. The Protein A conjugated chip was acquired from Xantec Ltd (Düsseldorf, Germany). A PBS solution with the addition of the 0.1% BSA and 0.005% Tween 20 was used as a running buffer. A 30 μg/mL RVT1 solution was loaded onto the chip and a series of the binding experiments with PF4 with the concentrations in range of 100-2000 nM were carried out. The acquired sensograms were corrected for the blank solution (matching the original solution of the PF4) and kinetics fitting was conducted in OriginPro software.

### Molecular modelling

The homology modeling of the VITT mAb Fab segment was carried out with a Schrödinger 2022-2 (Schrödinger LLC, New York, NY) suite using influenza A IgG1 (PBD 6URM) Fab as a template. MD simulations of the system using OPLS4 force field) were run for 250 ns after energy minimization and NVT equilibration for obtaining the representative structure of the RVT1 Fab. The VITT mAb Fab/PF4 docking was carried out with both Schrödinger-integrated PIPER algorithm and ZDOCK (56), yielding highly comparable results.

A detailed description of all experimental and modeling protocols is available in the ***Supplementary Methods*** section.

## Acknowledgments

This work was supported by a grant R01 GM112666 from the National Institute of Health. The authors are grateful to Prof. Leonid A. Pobezinsky and Prof. Barbara A. Osborn (UMass-Amherst) for helpful discussions.

## Supplementary Material for

**Figure S1.**
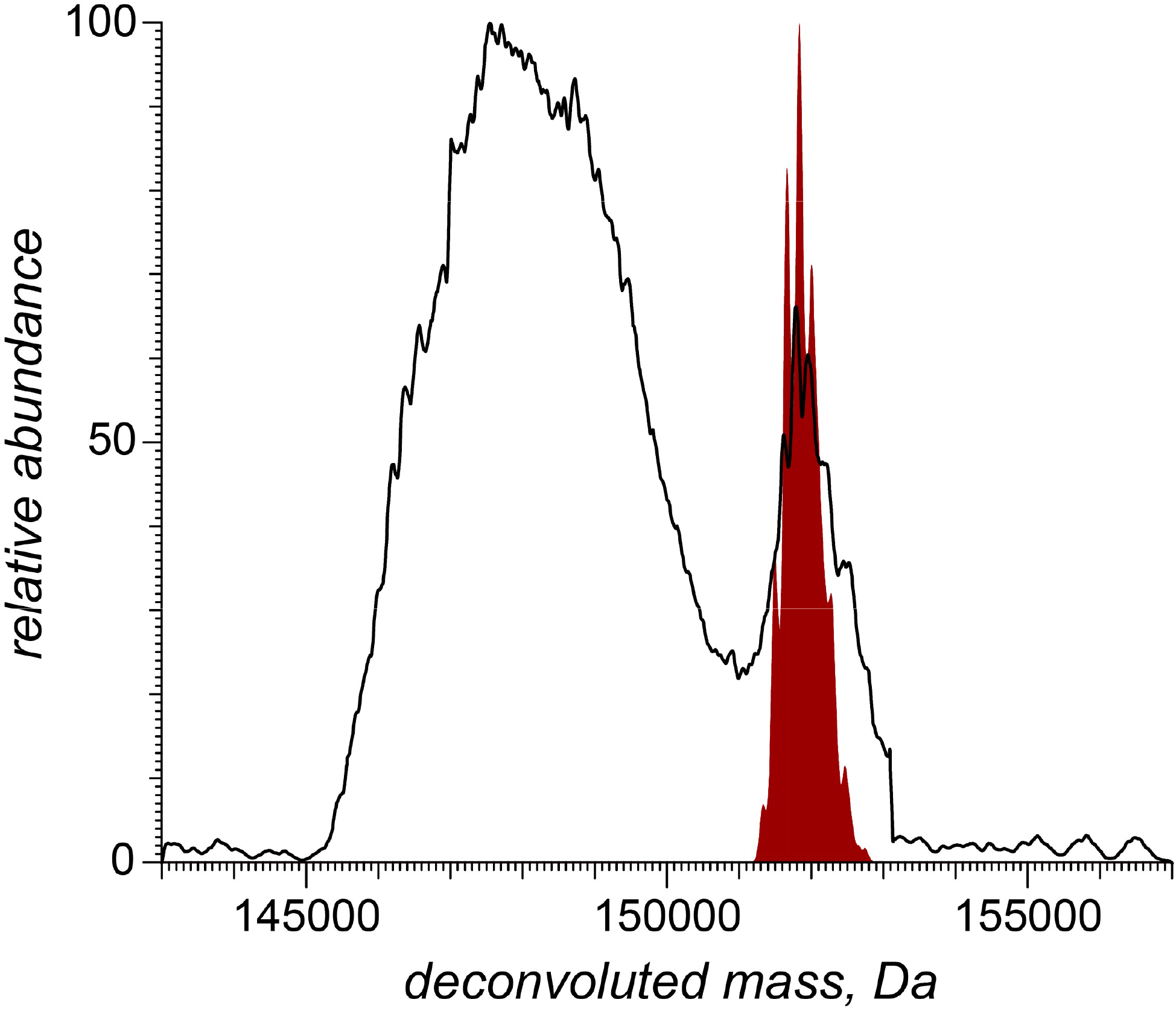
Mass distribution of anti-PF4 antibodies extracted from the VITT patient’s plasma. The distribution was obtained by deconvoluting the ESI mass spectrum of an aqueous solution of the antibodies in 150 mM ammonium acetate using a UniDEC algorithm. The mass and charge range selection for the decovolution process was made by supplementing MS measurements with limited charge reduction in the gas phase using a procedure describe elsewhere (2). The maroon trace represents the results of the mass convolution of the light chain and Fc/2 and Fd fragments of the heavy chain produced by IdeZ digestion and disulfide reduction of the entire antibody ensemble followed by LC-MS analysis (see Figure 1 in the main text).

**Figure S2.**
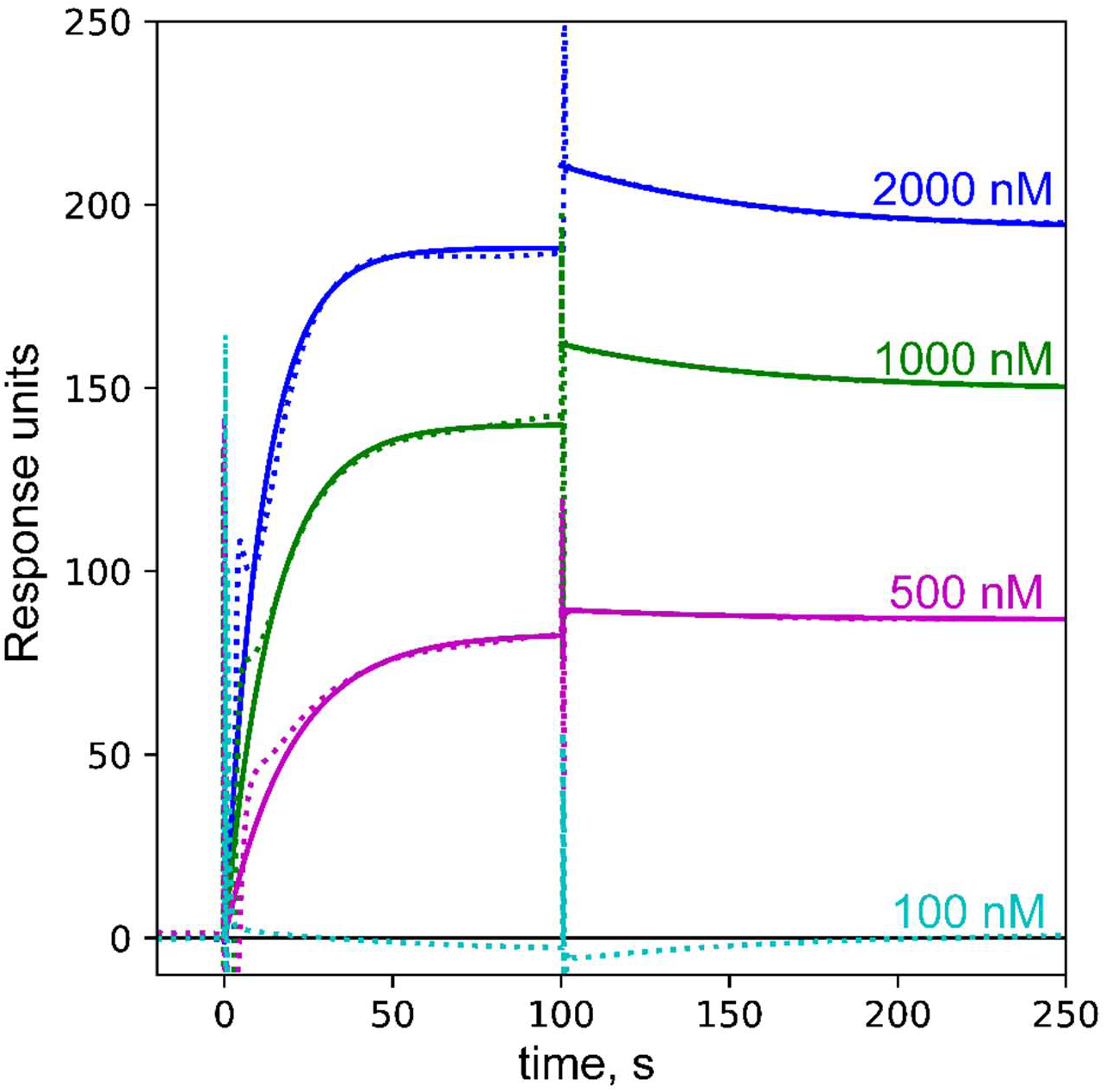
Surface-plasmon resonance sensograms obtained for the binding of the PF4 to the RVT1-captured Protein A chip. The concentration of the PF4 was titrated from 100nM till 2uM concentrations, and the signal was normalized to the sensograms with the blank solution whose composition matches the solution of PF4.

**Figure S3.**
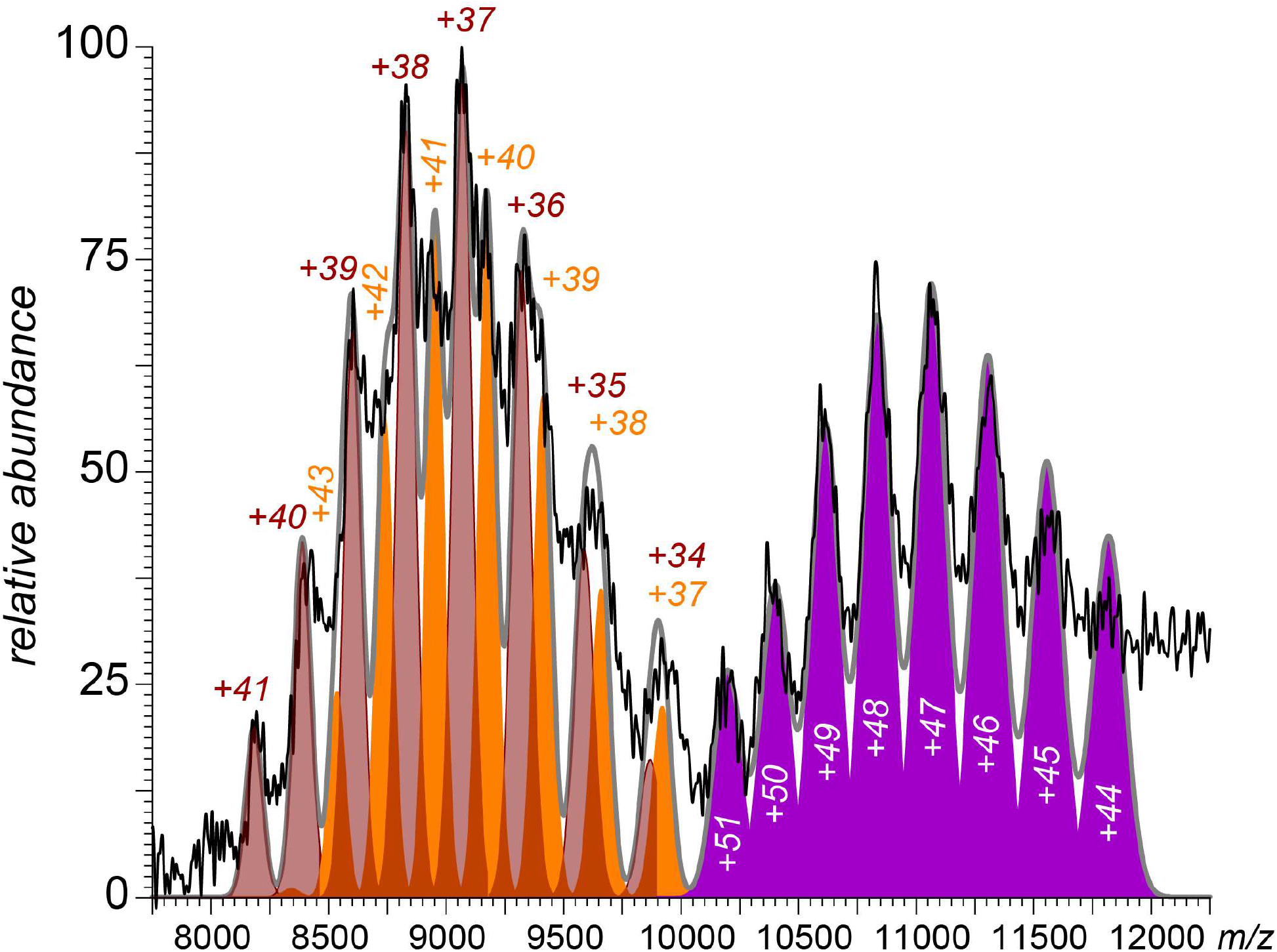
The native mass-spectra of RVT1:PF4 nearly-equimolar mixture fitted with mass-distributions of [2 RVT1+Fab] (brown), [2 RVT1+ 2 Fab] (orange) and [3 RVT1+ 2 Fab] (purple) at different charge states

**Figure S4.**
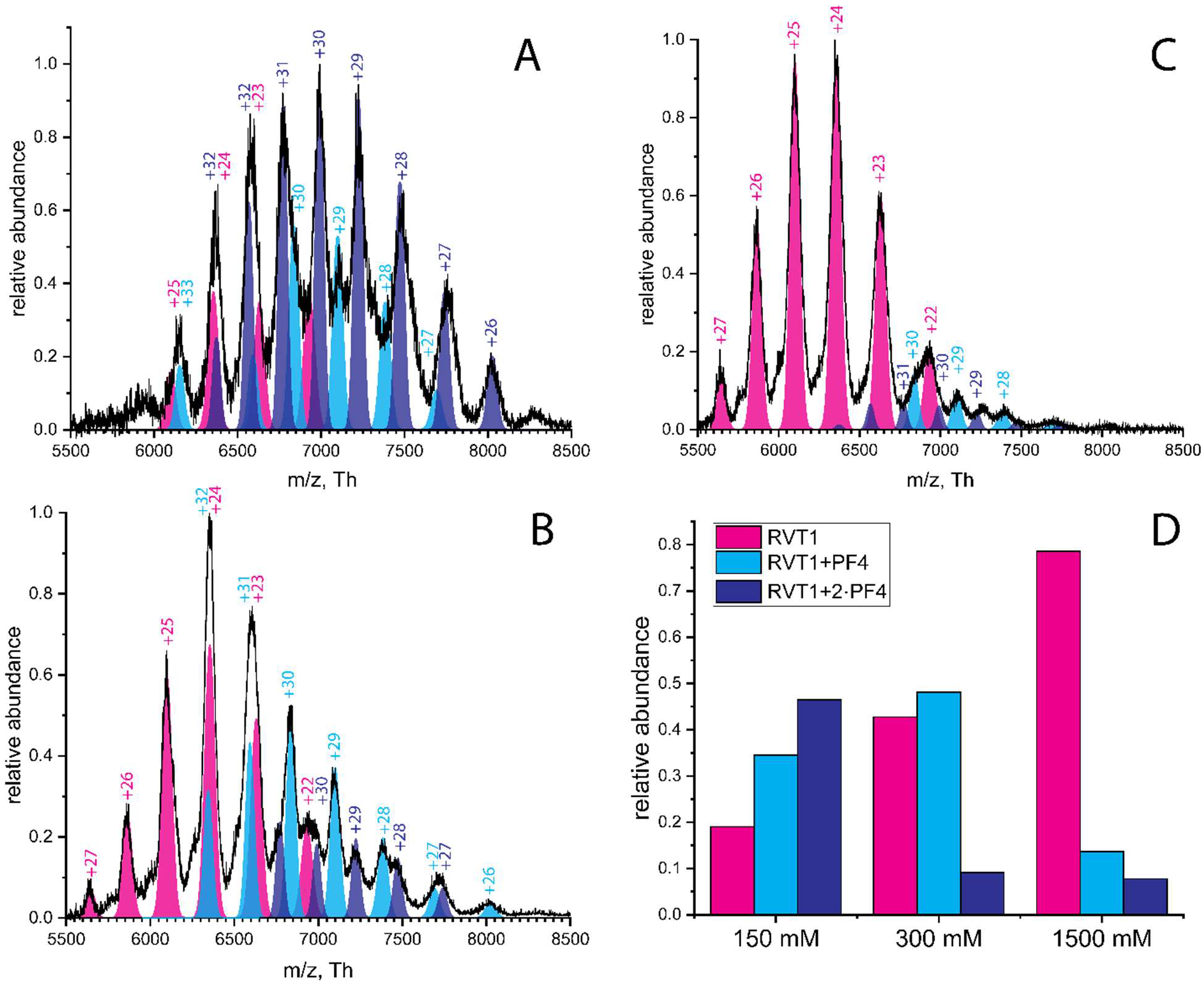
Native ESI mass spectra of a RVT1/PF4 (1:5 molar ratio) acquired at different levels of solution ionic strength.

**Figure S5A.**
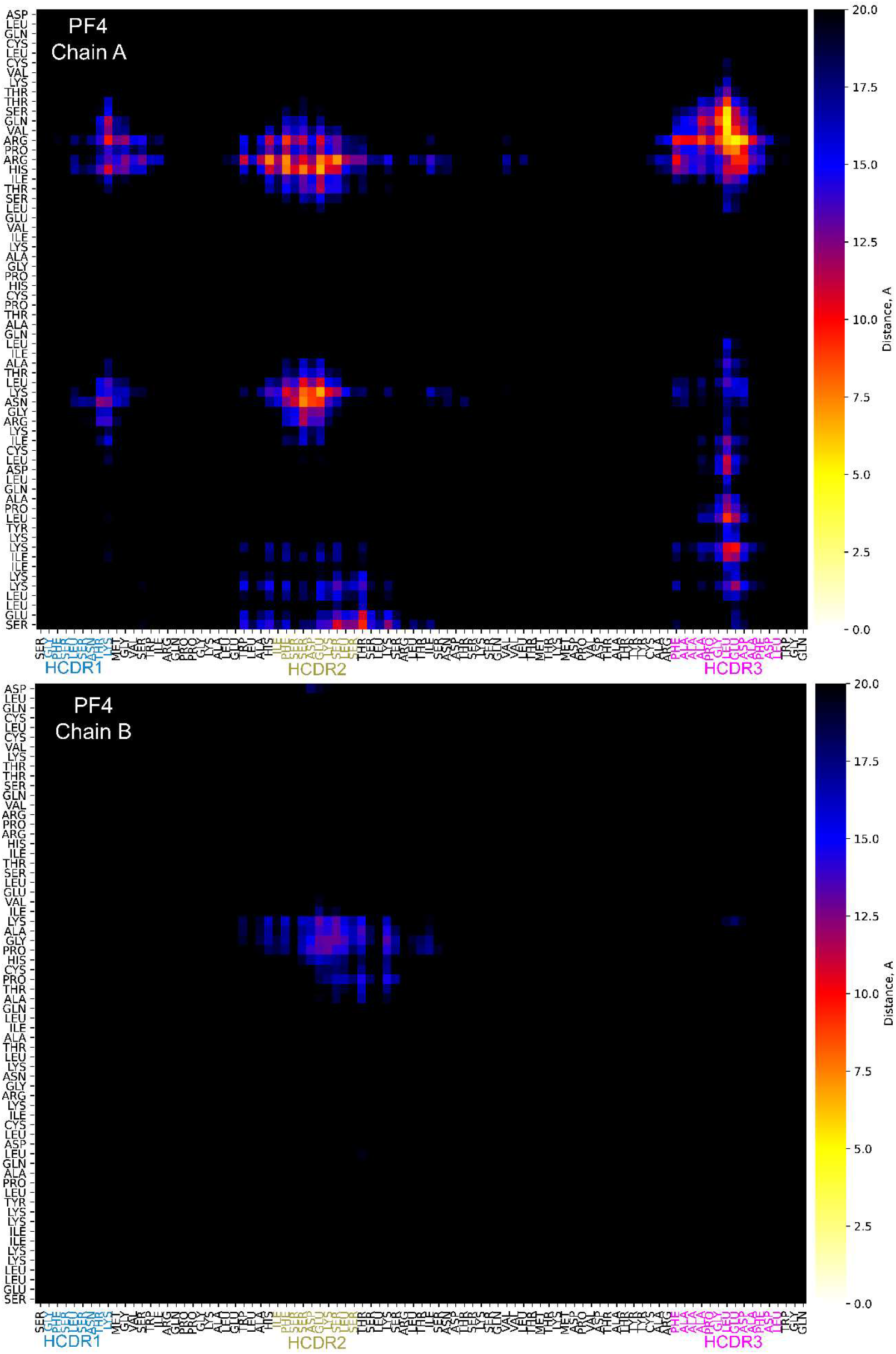
The distances between the RVT1 H-chain and the two nearest monomers of the

**Figure S5B.**
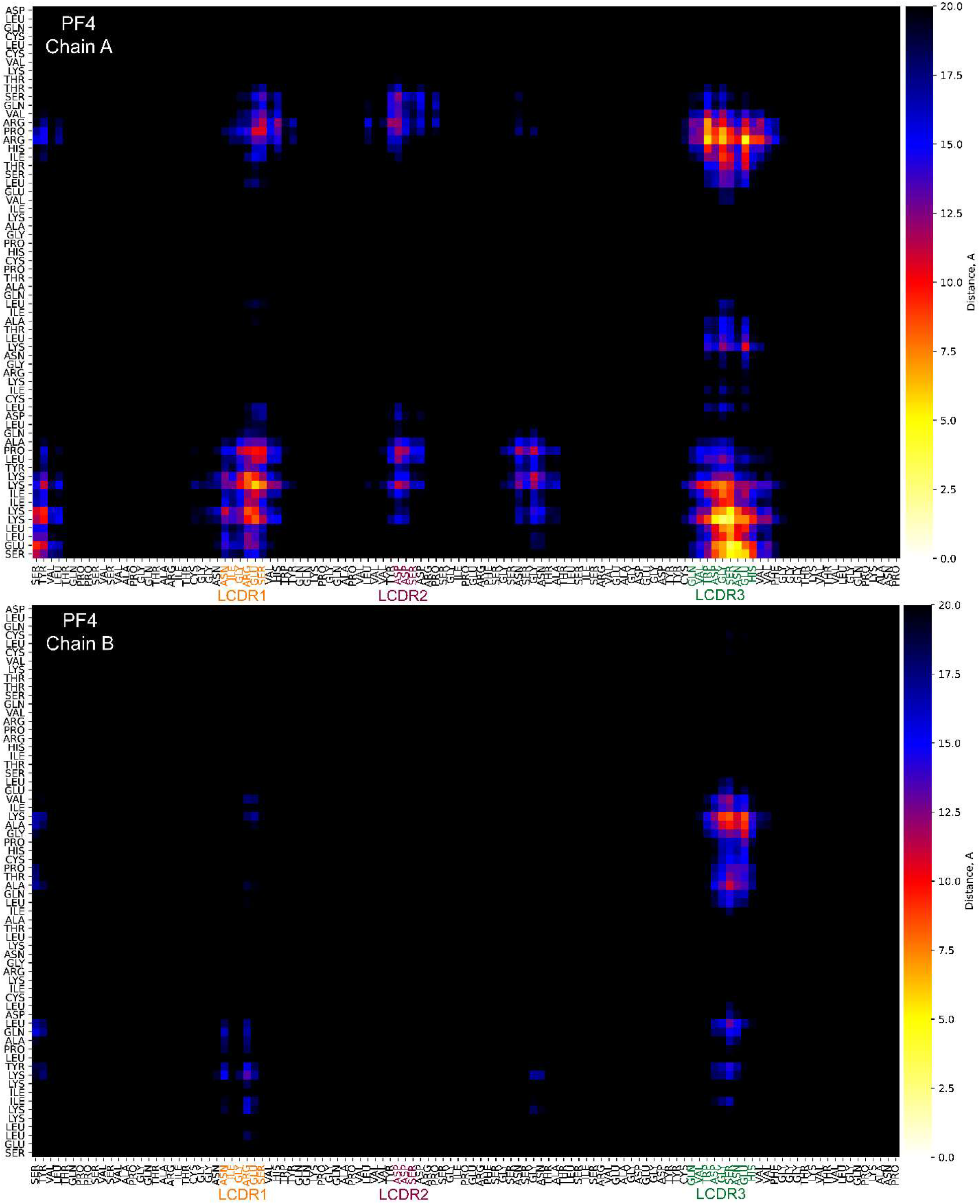
The distances between the RVT1 L-chain and the two nearest monomers of the

**Figure S6.**
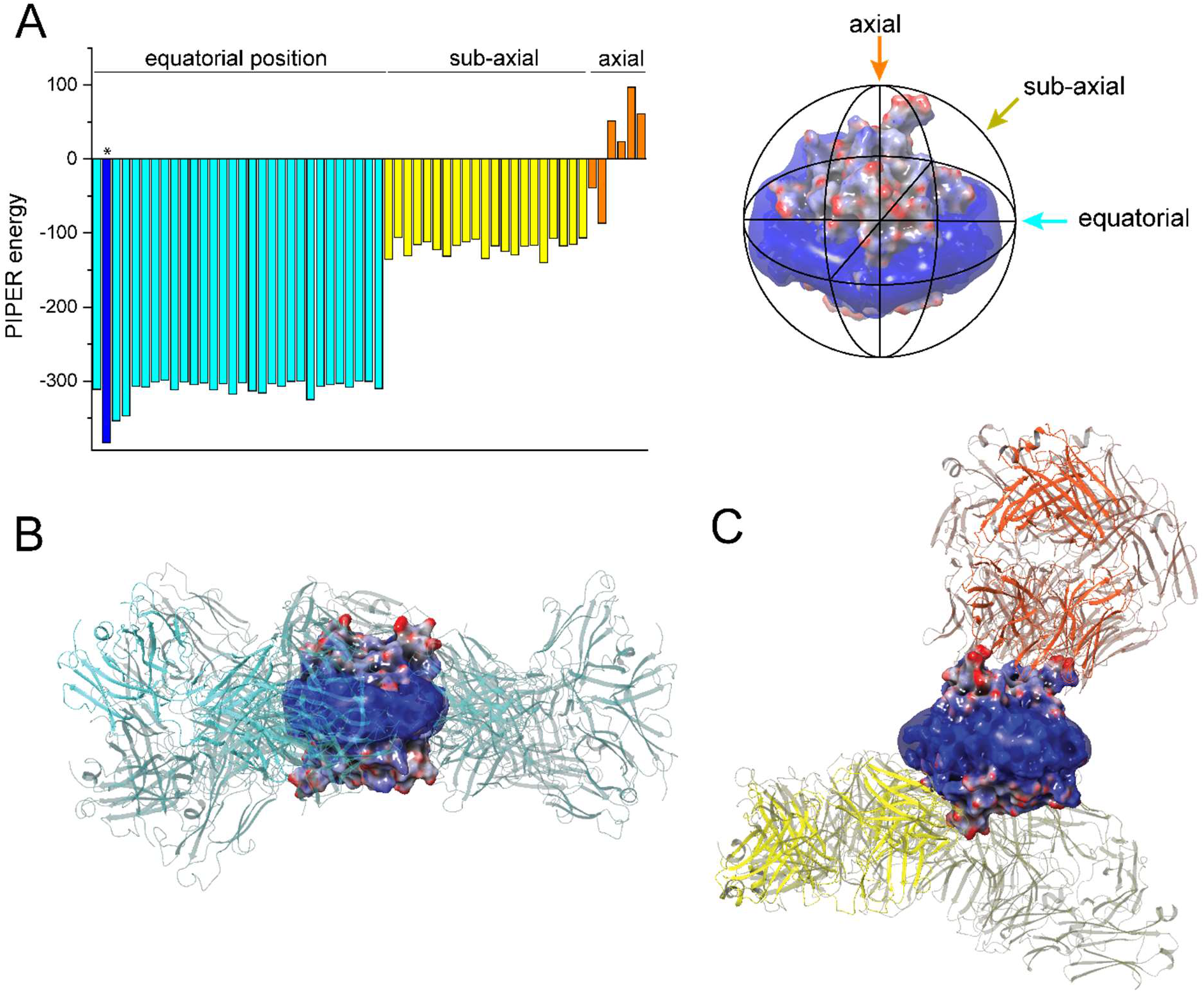
A. PIPER-based docking screening energies for RVT1 Fab docked to the various regions of PF4. Cyan columns refers to the docking energies after docking of RVT1 Fab to the equatorial belt of the PF4, yellow – to the sub-axial regions, orange – to the axial ones. The blue column with lowest energies refers to the docking of Fab to the equatorial belt of PF4 that matches epitope mapping data of VITT antibodies. **B.** Examples of the RVT1 Fab docked to the PF4 in equatorial positions. **C.** Examples of the RVT1 Fab docked to the PF4 in the sub-axial (yellow) or axial (orange) positions.

**Figure S7.**
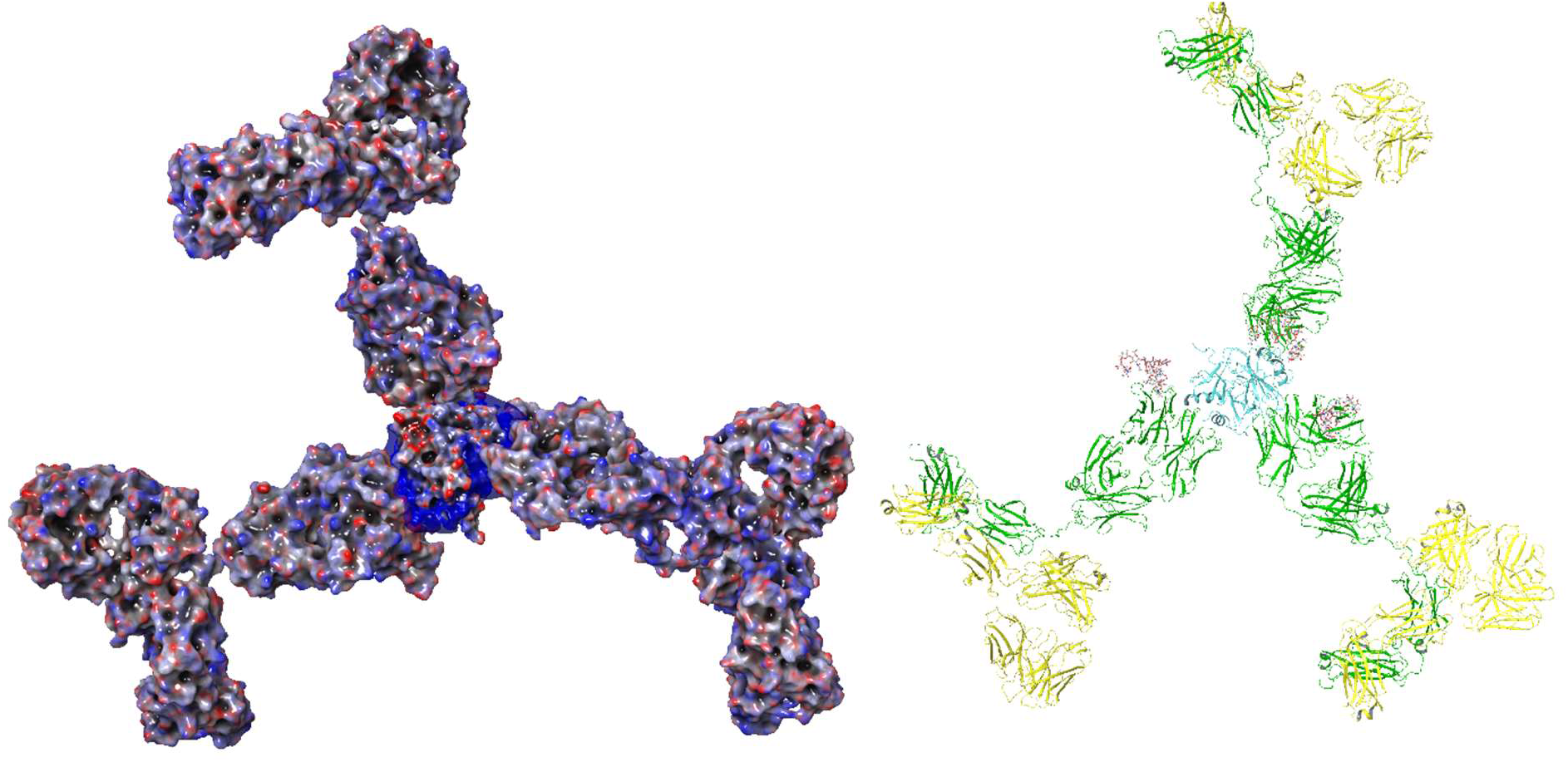
A dendritic mode of *RVT1* cross-linking by PF4 produced by molecular docking of three Fab segments to a single PF4 tetramer, followed by the extending the Fab fragments to full-length antibodies.

**Figure S8.**
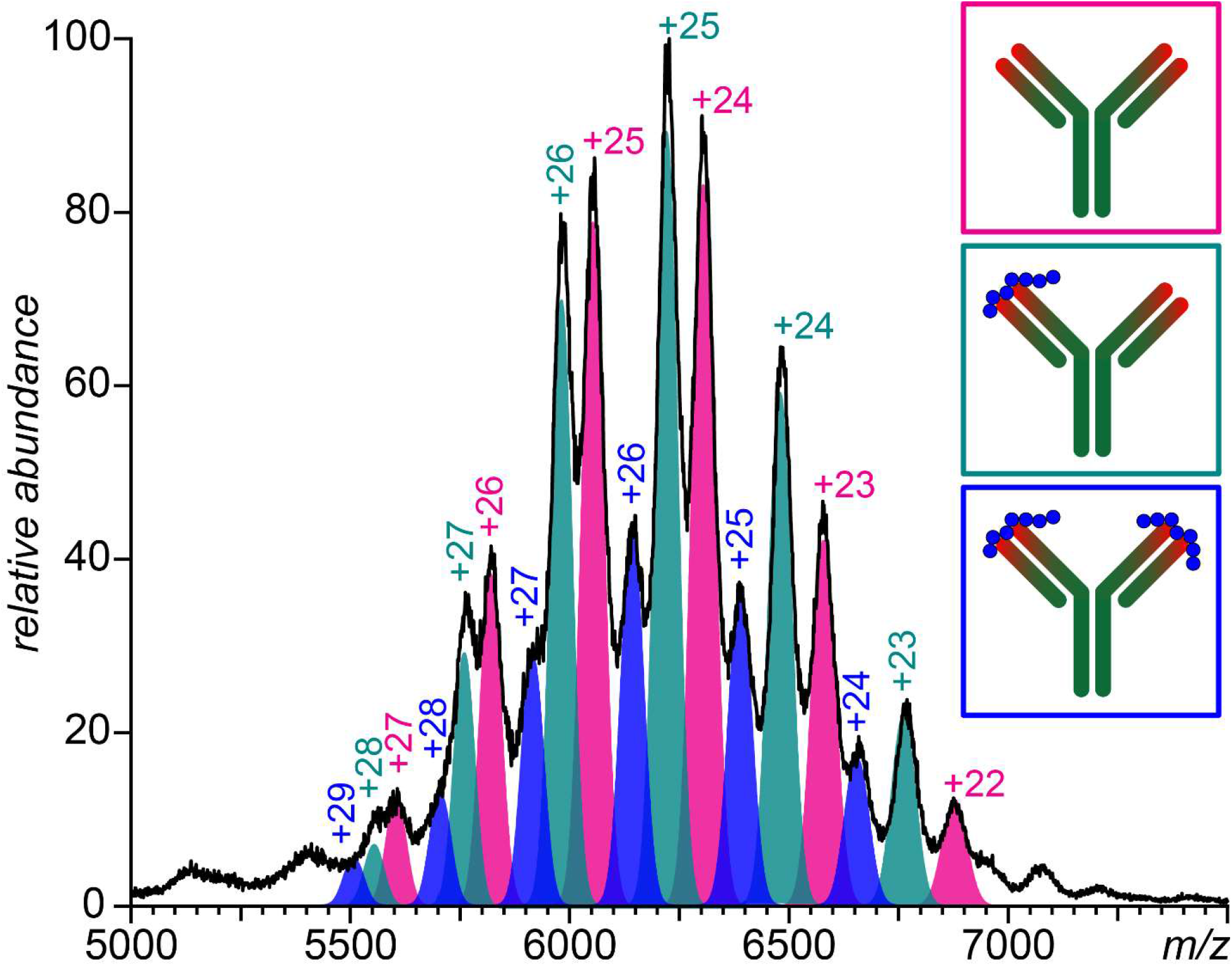
A native ESI mass spectrum of a 1:2 RVT1/protamine mixture acquired from an aqueous solution maintained at neutral pH and physiological ionic strength (150 mM ammonium acetate).

## Supplementary Methods for

### Study Participant Criteria

The sodium citrate plasma sample used for this study was obtained from a clinically diagnosed VITT patient referred for diagnostic testing to the McMaster Platelet Immunology Laboratory. Patient VITT diagnosis was based off the then-recent AstraZeneca vaccination, the presence of thrombotic complications and testing serologically positive for anti-PF4 IgG antibodies using the commercially available PF4-enhanced heparin-dependent IgG/A/M-specific EIA (Immucor), OD ≥ 0.45, and also positivity in the PF4-enhanced SRA (SRA ≥ 20% ^14^C-serotonin release). This study was approved by the Hamilton Integrated Research Ethics Board (HiREB).

### Anti-PF4 Antibody Purification

Anti-PF4 antibodies were purified from the patient plasma using the following protocol. Plasma was heated and treated at 56°C for 30 minutes and centrifuged at 21,000 g for 10 minutes to separate the sample supernatant. The isolated supernatant was diluted 4-fold with phosphate-buffered saline (PBS) and loaded onto a gravity flow column containing protein G sepharose beads (Protein G Sepharose 4 Fast Flow cat 17061801, Cytiva, Marlborough USA). The beads were washed with PBS until the flowthrough was observed to be <0.09 at 280nm. The bound antibodies were then eluted off the column using glycine (0.1 M, pH 2.8) and immediately neutralized using tris (2 M, pH 8.4). The eluted total IgG was buffer exchanged to PBS and concentrated to 7 mg/mL using 50 kDa centrifugal filter units (Ultracel-50, cat UFC905024, Amicon, Tullagreen IRL). The purified IgG fraction was incubated with biotinylated PF4 (0.1 mg/mL final concentration), generated in-house as previously described (1), for two hours at room temperature under gentle rocking conditions. Washed streptavidin sepharose beads (Streptavidin Sepharose High Performance, cat 17511301 Cytiva, Marlborough USA) were added to the PF4/antibody mixture (ratio of 1.5 mL 50% slurry to 2 mL antibody solution) and left to incubate overnight on a rocker at 4 ^O^C. The beads were then washed with PBS till O.D._280_ was <0.09. The bound antibody was then eluted off the beads as described before using glycine and tris. The eluted IgG was then mixed with 2 M NaCl at a 1:10 volume ratio and incubated for 15 minutes on a rocker. The sample was then centrifuged at 300 g, 1 minute and filtered through a 0.2 μm filter to remove beads before being buffer exchanged to PBS using a 50 kDa centrifugal filter. The final sample was then tested in the PF4-enhanced SRA to confirm reactivity.

### Intact-mass MS characterization of anti-PF4 antibodies

The enriched anti-PF4 antibodies (vide supra) were buffer exchanged using size-exclusion chromatography with SEC-3000 SEC column operated under the 1ml/min flowrate using the 150mM Ammonium Acetate, pH = 6.8 as a running solution. The eluted fraction of anti-PF4 was collected and concentrated using Vivaspin 500 MWCO 10kDa centrifugal filter (Sartorius). The final solution was loaded into the gold-coated borosilicate capillaries for the subsequent native mass-spectrometry analysis. Intact MS analysis was carried out using a Synapt G2 HDMS (Waters Corp., Milford, MA) hybrid quadrupole/time-of-flight mass spectrometer equipped with a NanoLockSpray nano-ESI ion source. The capillary voltage and nanoflow gas parameters were adjusted to establish a stable spray, and the sampling cone voltage was set to the maximum value (200 V) to ensure the efficient desolvation of ions.

### Intact-mass characterization of the large antibody fragments

The antibody segments LC, Fc/2 and Fd were generated by enzymatic digestion with IdeZ followed by chemical reduction of the disulfide bridges. The antibody was digested by the IdeZ (ThermoFisher, Waltham, MA) according to the standard protocol of manufacturer and then reduced by the 300 mM solution of DTT at 55 ^O^C for one hour in the dark. The fragments were separated using 4.6×100 mm, 3.5 μm pore size RP-mAb C4 column (Agilent Technologies, Santa Clara, CA) at a flowrate of 300 μL/min and room temperature. A SolariX 7 (Bruker, Billerica, MA) Fourier-transform ion cyclotron resonance mass spectrometer was used for on-line detection of the large proteolytic fragments.

### Peptide mapping and de novo-sequencing of the antibody

The antibody peptide mapping was carried out independently with either trypsin or chymotrypsin (New England Biolabs, Ipswich, MA), and the fragment peptides were analyzed with LC-MS/MS. The on-line separation was carried out with an Easy-nLC 1000 (Thermo, San Jose, CA) system using a Thermo PepMax Acclaim C18 column. The on-line detection and MS/MS analysis of the peptides were performed with an Orbitrap Fusion (ThermoFisher, Waltham, MA) mass spectrometer equipped with a NanoSpray Flex ion source. Ion fragmentation in the gas phase was carried out with HCD, and the annotation of fragment ions was done with Peaks Studio Xpro, followed by the manual assembling of the protein sequence. Further details of the data acquisition and interpretation processes are available in a technical report deposited to bioRxiv (2).

### Functional Platelet Activation Assays

Platelet-rich plasma was prepared by differential centrifugation of whole blood (250 × g 10 min) collected using acid-citrate-dextrose (ACD) from pre-screened donors. Platelets were radiolabeled with ^14^C-serotonin and then washed twice in calcium/albumin-free Tyrode’s solution (CAF) and resuspended in albumin-free Tyrode’s buffer (AFT) at 350,000 platelet/µL. The platelets were then incubated for 60 min at 22 ^O^C with RVT1 (8 µg/mL). For the regular SRA, pharmacologic concentrations of unfractionated heparin at concentrations of 0, 0.1, 0.3 and 100 U/mL were added during this incubation step. For the PF4-enhanced SRA the heparin was replaced with exogenous recombinant PF4 (generated in house) and added at 0, 10, 25 and 50 µg/mL concentrations (3). The reactions for either assay were then stopped using 5 mM EDTA in PBS, and the supernatant of each reaction was removed and counted in a scintillation counter (Packard Bell, Topcount) to measure the release of ^14^C-serotonin. The amount of serotonin released in each well was compared to the total radioactivity in the platelet preparation and expressed as a percentage of the total amount used. The SRA result was considered positive if the sample caused ≥20% serotonin release in either assay (4). In addition to test sera, positive and negative control sera from established samples were used to confirm assay functionality. In some conditions, platelets were also preincubated with the monoclonal anti-CD32 antibody IV.3 (5) to confirm activation was FcRγIIa dependent. All conditions were tested in technical duplicates, and experiments were repeated to confirm the findings.

### Surface-plasmon resonance (SPR) analysis

Surface-plasmon resonance analysis was carried out using a Biacore T200 (Cytiva) device. A protein A conjugated chip was acquired from Xantec Ltd (Düsseldorf, Germany). As a running buffer, the solution of PBS with the addition of the 0.1% BSA and 0.005% Tween 20 was used for the removal of non-specific binders. A 30 μg/mL RVT1 solution was loaded onto the chip and a series of binding experiments with PF4 in the concentration range 100-2,000 nM were carried out. The acquired sensograms were background-corrected using a blank solution (that matches the original solution of the PF4) and kinetics fitting was conducted in OriginPro software.

### Native MS analysis using a Synapt G2 mass spectrometer

Native MS measurements of RVT1 interactions with recombinant PF4 or protamine were carried out after buffer-exchanging the proteins with a Superdex 75 column into the 150 mM ammonium acetate solution, pH 6.8. The protein fractions were concentrated using Amicon Ultra-4 concentration cartridges with different MWCO (3 kDa for PF4 and 10 kDa for RVT1). A Synapt G2 HDMS (Waters Corp., Milford MA) hybrid quadrupole/time-of-flight mass spectrometer was used in the nano-ESI mode for native MS data acquisition. To enhance the transmission of the high-MW complexes, the backing pressure was increased up to 7 mBar. The sampling cone voltage was optimized to enhance the desolvation of ions while keeping the non-covalent complexes intact. The acquired data was analyzed using MassLynx 4.2 or OriginPro software.

### Homology modeling of RVT1 Fab

Molecular modeling work was carried out with a Maestro platform (Schrödinger Release 2022-2, Schrödinger LLC, New York, NY) using OPLS4 force field. The RVT1 Fab segment model was built by homology modeling based on the crystal structure of human anti-influenza A IgG1 (6URM). All modeled systems were neutralized with appropriate amounts of Na^+^ ion and Clˉ ions to achieve a total salt concentration of 150 mM. The final solvated box contained ca. 87,000 SPC water molecules and its size 180×90×200 Å was sufficiently large to prevent proteins contacts with their periodic images. The production simulations were run at 310 K for 250 ns. A Noose-Hoover thermostat was used to control the temperature and the 1 atm pressure was maintained with a barostat (volume move attempt every 2 ps). All bond lengths involving hydrogen atoms were constrained using the SHAKE algorithm to allow for an integration time step of 2 fs with the RESPA algorithm. Long-range electrostatic interactions were treated using the particle mesh Ewald method, and the short-range van der Waals interactions were treated with a 9 Å cutoff.

### Molecular docking of Fab to PF4 units

Fab/PF4 docking studies were carried out independently using (a) the protein-protein docking function in Maestro and (b) ZDOCK. The Maestro-based docking was carried out with PIPER by sequential docking of two RVT1 Fab segments to a single PF4 molecule by selecting the highest-ranking structures followed by a short energy minimization and equilibration step. For Maestro-based docking the PF4 structure was built using 4R9Y as a template, to which the six missing N-terminal residues were appended followed by energy minimization. MD simulations of the PF4-Fab complexes were carried out after equilibrating the initial constructs produced by docking using two rounds (100 and 12 ps) of NVT simulations by slow heating from 10 to 310 K, while restraining all heavy atoms using harmonic potentials. This was followed by a 24 ps-long simulation round at 310 K with no restraints on the heavy atoms. For ZDOCK-based docking, the intact 4R9Y structure was used for sequential docking, and no equilibration steps were performed between the docking of protein units. The contact maps were generated using an in-house developed Python-based script.

Monitoring the antigen-antibody docking at different sites was carried out with PIPER by masking non-CDR region, *i.e*. the attractive potentials for residues in the non-CDR region were removed. The Chothia definition was used to define the CDR region. Three different criteria were used to acquire different docking poses. For equatorial docking, docking was done with no restrains. For sub-axial docking, distance restraints were introduced between E100 residue of HCDR3 of the antibody and A2 in single chain of PF4 tetramer. For axial docking distance restraints were introduced between E100 in HCDR3 and single chain of PF4 tetramer, and between N95 in LCDR3 of the antibody and A2 of the opposite single chain of PF4 tetramer. The distance restraints are mainly introduced to confine the relative position of PF4 tetramer and the RVT1 Fab and the minimum and maximum distance between the two selected residues is set at 2 and 10 Å, respectively.

